# Impact of ectopic calcification on vascular oxygenation in placental dysfunction models: Simultaneous *in vivo* assessment with multi-parametric ultrasound and photoacoustic imaging

**DOI:** 10.1101/2023.12.04.569997

**Authors:** Skye A. Edwards, Ana Correia-Branco, Deeksha M. Sankepalle, Aayush Arora, Allison Sweeney, Patrick R. Solomon, Mary C. Wallingford, Srivalleesha Mallidi

## Abstract

Preeclampsia (PE) is a prevalent gestational disorder occurring in 2-10% of all pregnancies. It is associated with placental dysfunction which can result in maternal mortality and preterm birth. Despite scientific advances, the underlying mechanisms of PE and the progression of placental dysfunction remain poorly understood without preventative or curative treatment. Other than blood tests and postdelivery placental diagnostics, there are limited detection methods for monitoring placental health. A major roadblock is the lack of imaging modalities available to monitor placental hemodynamics, as the commonly used clinical imaging systems are either harmful due to ionizing sources, too expensive and inaccessible, or provide insufficient data. We propose the combination of quantitative ultrasound and photoacoustic imaging (QUSPAI) to characterize the material and structural changes of the placenta during pregnancy in normal and disease states. The Slc20a2 murine model of placental dysfunction has previously been characterized by increased ectopic placental vascular calcification, reduced fetal growth, and decreased postnatal bone mineral density. In this study we utilized the Slc20a2 model and imaged knockout (KO) and wildtype (WT) placentae on embryonic day (E)18.5. The dual-wavelength images and radio frequency data were collected *in vivo* and *ex vivo* to provide quantitative ultrasound spectral (QUS) parameters and blood oxygen saturation. The photoacoustic images indicated that the WT tissue had statistically higher blood oxygen saturation (StO_2_) values than the KO tissues while QUS data revealed strain-specific calcification effects. Building upon the E18.5 gestation data collected here, future work will involve monitoring longitudinal changes in these parameters, including early calcification, to support translating this method for early diagnosis of placental vascular diseases.

## Introduction

The placenta is a temporary organ that develops during pregnancy and serves a central role in facilitating the transportation and transfer of gas, nutrients, waste, and hormones between the maternal and fetal blood streams^1, 2^. Three critical categories of placental health can be monitored, including anatomical structure, metabolism, and vascular function. Placental dysfunction and insufficiency can impact each of these categories and cause numerous issues such as fetal growth restriction (small size for gestational age), spontaneous pregnancy loss, or preterm birth.^3^ Placental insufficiency is also observed in association with preeclampsia (PE), which affects 2-10% of pregnant individuals worldwide and being one of the top causes of the maternal deaths in the United States each year.^4-7^ Despite this fact, very little is known about PE and there are no early symptoms or curative treatments for PE, with the exception of the delivery of the placenta which can necessitate premature birth of the baby.

There are several imaging modalities that are used clinically and preclinically to probe into placenta and fetal development.^2^ Optical approaches such as confocal microscopy and optical coherence tomography offer high spatial resolution, however these modalities have poor penetrative depth (on the order of few microns to a few millimeters for confocal microscopy and optical coherence tomography respectively).^2, 8^ The common clinical imaging techniques, along with their respective modality variants, such as magnetic resonance imaging, computed tomography, positron emission tomography, and single photon emission computed tomography, have better penetrative depth, but poor spatial resolution, and they can be ionizing or require exogenous contrast agents that can be harmful to a pregnancy.^2, 9^ A modality that is non-ionizing, non-invasive, has decent resolution and penetration depth while being low cost and mobile is ultrasound (US) imaging and hence is ubiquitously used in evaluating placental health. Indeed, the Grannum grading system utilizes US images during routine pregnancy checkups.^10-12^ The grading system was established in 1979 and relies on an observer to determine the presence of calcification in the placenta based on the echogenicity of the US image.^12^ This method of placental grading via qualitative US has high variation between in intra-observer and inter-observer diagnoses and reflects the need for specialized training.^13^ Efforts have been made to use many different imaging modalities to investigate the placenta’s structure and function from the nano to macro scale.^2, 9, 14, 15^

The gold standard of placental and fetal imaging, namely US imaging, could aid in assessing differences in placenta health when supplemented with quantitative methodologies such as analyzing the ultrasonic radio frequency (RF) echoes to estimate physical tissue properties such as acoustic scattering.^16, 17^ Quantitative ultrasound analysis investigates the normalized Fourier transform of user specified RF windows. The scattering behaviors and estimated properties are associated with several parameters such as the mid-band fit (MBF), zero-slope intercept (ZSI), and spectral slope (SS). The effective scatterer size impacts the SS, ZSI, and MBF. The concentration of scatterers influences the ZSI and the MBF. Lastly the acoustic impedance only determines the MBF.^17, 18^ It is shown that there are statistical increases in the spectral parameters in pre-treated and post-treated tumors due to the apoptosis.^19-23^ QUS has also been employed in the understanding of nonalcoholic fatty liver disease (NAFLD) where the higher lipid contents in patients with NAFLD are less homogeneous and have larger spectral parameters.^24, 25^ Other image features and ratio metrics can be added into the spectral analysis post-data acquisition to further quantify anomalies in a range of tissues.^21, 26^ Applying a quantitative method that utilizes the raw ultrasonic RF data can provide insightful information and has the potential to support the placental calcification grading system.

The newer emerging modality, namely photoacoustic (PA) imaging, is a promising modality to image vascular structure and function of placentae in a safe and efficient manner. More importantly, PA imaging is a non-ionizing modality that can be combined with US seamlessly as they share the same receiver electronics.^2^ Specifically, PA imaging is based on generation of photoacoustic signals via thermoelastic expansion and contraction of the absorber in tissue after irradiation with a nanosecond pulsed laser. The generated PA signal is a function of the input laser irradiation wavelength and represents the optical absorption properties of the absorber. For example, blood (hemoglobin) is a major absorber in the body that provides good photoacoustic contrast. The oxygenated and deoxygenated hemoglobin chromophores having distinct optical absorption properties enabling calculation of blood oxygen saturation (StO_2_) calculated as ratio of oxygenated hemoglobin to total hemoglobin (HbT). Performing multi-wavelength PA imaging of the tissue can aid in deciphering the 3D map of StO_2_ in the placenta. Furthermore, PA imaging can be combined with US imaging, and without need of additional receiver electronics and image co-registration, structural and functional information on the placenta can be obtained. For example human placentae, have been imaged *ex vivo* with US and PA imaging termed, USPA, after a cesarean section and these studies have demonstrated that the acoustic spatial resolution is sufficient to provide insights to the structure and function of the placenta.^27, 28^ The current limitation in light penetration depth through the human body only allowed previous studies to investigate the function of the placenta at the time of delivery and not throughout the gestational period. Furthermore, due to this depth limitation, in the past several years, most *in vivo* imaging studies involving longitudinal measurement of placental oxygenation under normal, hypoxic or hypertensive conditions have been performed in small animal rodent models^29-36^ while neonatal large animal (e.g., piglets) were used for assessing StO_2_ in the brain sagittal sinus.^37^

Previous placental dysfunction studies have utilized a variety of animal models ranging from rodents to non-human primates. The models that were investigated in most of the studies required the induction of placental dysfunction via drug administration, surgical implantation, or artificial environment.^29-36^ Reduced uteroplacental perfusion pressure (RUPP) rat models are among the most common placental ischemic models.^38^ This model is often used to investigate the uterine perfusion pressure, the resultant placental morphology, and potential treatments, but requires survival surgery to induce the model.^30, 38, 39^ Only one study researching placental response to an oxygen challenge utilized a genetically modified hypertensive mouse model to compare to the control and drug induced model.^33^ The sodium dependent phosphate transporter gene Slc20a2 is expressed in the human and mouse placenta and it protects against age-associated vascular calcification.^10, 40^ Altered Slc20a2 expression is associated with placental dysfunction, preeclampsia, fetal growth restriction, and placental calcification associated with thickened basement membranes.^40^ The Slc20a2 genetic knockout (KO) model, allows for an altered placenta and restricted fetal growth without the need for surgical intervention.^40^ In this study, we hypothesize that hyper calcification will impact the function of the placenta in the late gestational periods. We established placental calcification and altered function on embryonic day E18.5 in the Slc20a2 deficient mouse model. Using StO_2_ along with ZSI and MBF parameters, differences in oxygenation and calcification levels were studied across the two placental genotypes.

## Materials and Methods

### Animal Studies

All animal studies were conducted under the appropriate protocols approved by the Tufts University Institutional Animal Care and Use Committee. Wildtype (*Slc20a2*^+/+^) C57Bl/6J mice (Jackson Labs) and C57BL/6NTac-Slc20a2^tm1a(EUCOMM)Wtsi^/Ieg (*Slc20a2* KO-first +/−) mice (European Mouse Mutant Archive) were purchased, and *Slc20a2*^+/-^ KO-first were intercrossed to establish the colonies, mice, and placentae used throughout the experiments.^40^ At least 3 maternal models for both (*Slc20a2*^+/+^) and (*Slc20a2*^+/-^) were analyzed. Twelve hours prior to imaging the animals’ abdomen, fur was clipped down to make depilation easier and avoid chemical burns from depilation cream. As demonstrated in Fig. 1, the animals were imaged under anesthesia on gestational day E18.5 under 1-1.5% isoflurane. Immediately after the USPA imaging scan, pimonidazole hydrochloride was administered at 60 mg/kg via tail vein injection, one hour after injection the mice were euthanized, and the placentae and yolk sac of each embryo were extracted. The extracted placentae were imaged *ex vivo* placed in Phosphate Buffered Saline (PBS). Post imaging, the placentae were placed in formalin or O.C.T. Compound (Fisher HealthCare) for immunohistochemistry. The yolk sac was placed into Eppendorf tubes and placed on ice and then stored in a -20 °C freezer to be used for determining the respective placentae genotype.

**Figure 1:**
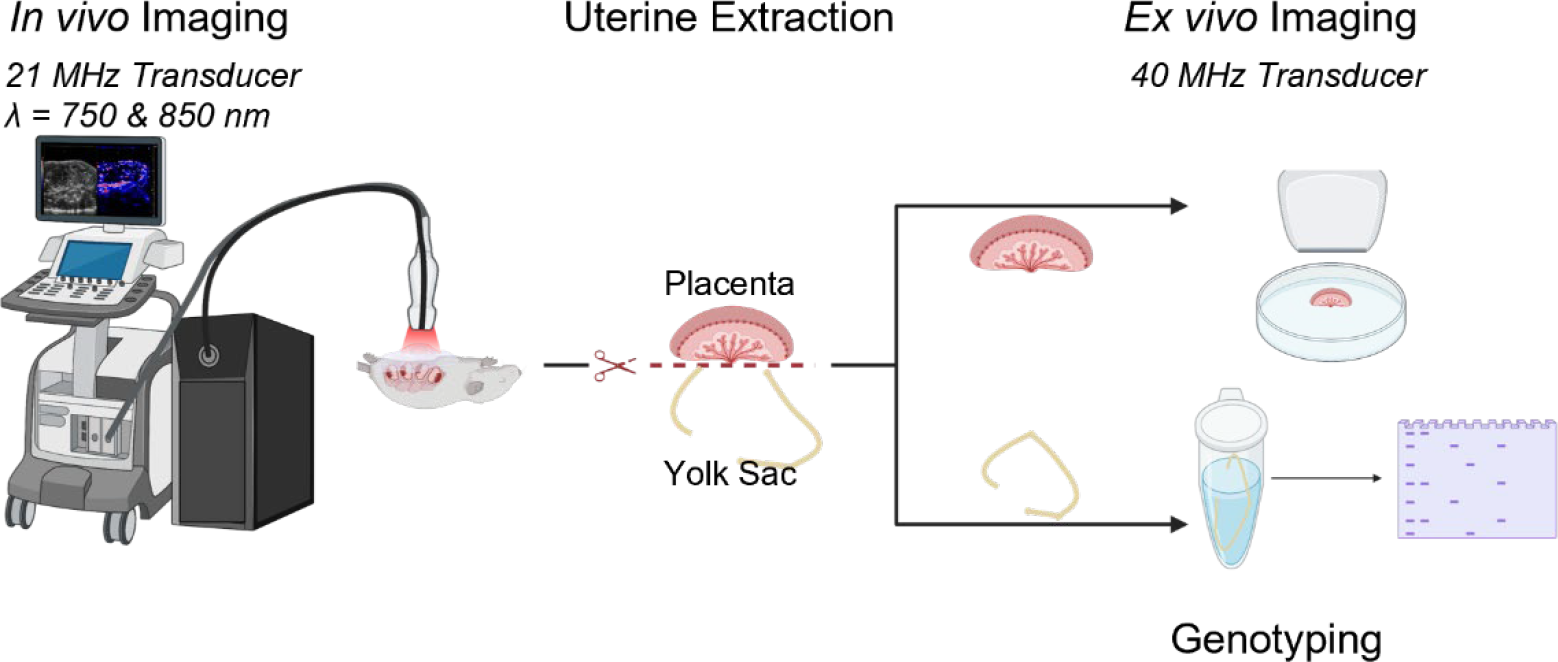
Schematic representation of the experiment workflow. Pregnant mice at gestational age E18.5 were anesthetized and placed in supine position. US imaging was performed to locate the placenta followed by PA imaging. Immediately after euthanasia, *ex vivo* imaging of the placenta was performed using a high frequency transducer and yolk sacs were collected to obtain the embryo genotype.

### DNA Extraction Method

Each extracted yolk sac was lysed with a mix 50 μL of tail buffer and 0.75 μL of proteinase K (Sigma-Aldrich). The samples were then incubated at 55 °C for at least one hour until the sample has degraded and then incubated at 95 °C for 10 minutes to deactivate the proteinase K. The DNA concentration was measured with a NanoDrop One (Thermo Scientific) and diluted to a concentration of 50 ng/μL. The diluted sample (4 μL) was then combined with 21 μL of polymerase chain reaction reagents. The mixture included 1 μL of custom primers, 12.5 μL of GoTaq Green Master Mix 2x (Promega), and 5.5 μL of Ultrapure DNase/RNASE-Free Distilled Water (ThermoFisher Scientific). Each sample was then processed in a thermo cycler with custom cycle times, the 10 μL of the amplified DNA samples were evaluated by agarose gel electrophoresis, which was ran for 50 minutes. The gel was prepped with 1x Tris-acetate-EDTA and 5 μL of SYBR safe (company Name). Once the gel electrophoresis finished, the gel was imaged with a FluorChem (ProteinSimple) in the SYBR Green setting.

### Photoacoustic and Ultrasound Imaging

Image acquisition was performed using the Vevo LAZR-X USPA imaging system (Fujifilm, VisualSonics, Inc.) equipped with a nanosecond OPO pulsed laser. *In vivo* USPA images were collected at 750 and 850 nm wavelengths and a 21 MHz transducer (MX250S). The PA and US gain were set at 45 and 22 dB, respectively. US was first utilized to approximate the number and location of visible embryos and placentae. 3D RF scans were performed on each visible embryo and placenta pair, followed by 3D US-PA scans of 1-2 superficial placentae. *Ex vivo* RF scans were conducted with a 40 MHz transducer (MX550) with a gain of 20 dB. The 3D StO_2_ and HbT maps were exported through the Vevo LAB Software (FujiFilm, VisualSonics, Inc.). Both maps were fluence compensated and the averages which do not include zero values, which ensures that areas with no blood vessel signals are not included in the overall value for each frame, were calculated for each placenta. RF scans of reference phantom made with known sizes of glass beads were collected routinely to determine the reference spectrum used for normalization^16^

### Fluence Compensation

The *in vivo* PA images were compensated for depth-dependent fluence attenuation utilizing Monte Carlo (MC) simulations of light propagation. Briefly, the 3D US images were segmented into 5 regions of interest (background, skin, embryo, placenta, and standard tissue). Each region was assigned the optical properties provided in Table 1. The size of the simulation geometry was determined by each image size with a bin size of 0.1 mm. The simulations were performed using Monte Carlo Extreme at 750 nm and 850 nm with 5 million photons launched from each bifurcated optical fiber.^41^ The resultant fluence maps were normalized and applied to the non-background regions of the PA image.

**Table 1.** Optical Properties used to fluence compensate PA scan of *in vivo* Placenta.

| Tissue | $\mu_a$ (mm<sup>-1</sup>)<br>750 nm \| 850 nm | $\mu_s$ (mm<sup>-1</sup>)<br>750 nm \| 850 nm | Anisotropy | Refractive Index |
| --- | --- | --- | --- | --- |
| Skin | 0.2504 \| 0.2935 | 25.22 \| 21.90 | 0.9 | 1.37 |
| **Placenta (liver)** | 0.2083 \| 0.2729 | 8.486 \| 6.911 | 0.9 | 1.37 |
| **Fetus (bone)** | 0.0012 \| 0.0020 | 13.18 \| 12.63 | 0.9 | 1.37 |
| **Standard Tissue** | 0.0110 \| 0.0199 | 10.97 \| 9.418 | 0.9 | 1.37 |

### Quantitative Ultrasound Processing

The RF data was processed, as seen in Fig. 2, by calculating a power spectrum for sliding windows across the placental region of interest (ROI). The sliding windows had a 95% overlap in the axial direction. Spectral parameters were determined by applying a linear regression between the bandwidth of the transducer for each sliding window. The frequency of the bandwidth was determined by utilizing a carbon fiber strand or a glass bead phantom, then taking a B-mode image, and analyzing the RF power spectrum at full-width half-max. For the 21 MHz transducer used, the bandwidth was 6.0669 – 33.2808 MHz and for the 40 MHz transducer the bandwidth was 9.5833-48.3438 MHz, averaged across six carbon fiber and glass beads images. The parameters MBF, ZSI, and SS were taken from the linear fit of each sliding window across the entire ROI, which was normalized by the power spectrum obtained from the reference phantoms, creating heatmaps for each 2D frame.^16^ The 2D frames were then segmented on MATLAB, combined into tiff images, and uploaded onto AMIRA (ThermoFisher Scientific) to create 3D reconstructions of the *in vivo* and *ex vivo* visualizations. Parameter ratios were determined by thresholding all the collected frames by a range for ZSI and MBF, 50 dB and 40dB, respectively. The number of pixels above the threshold per the number of US pixels between 40-80 dB, were then used to determine the ZSI and MBF ratios. Area percentage was specifically calculated by taking the number of MBF or ZSI pixels above the determined threshold and divided by the number of US pixels present in the frame. Then each frame had an area percent value, which is the averaged for each placenta in each genotype.

**Figure 2:**
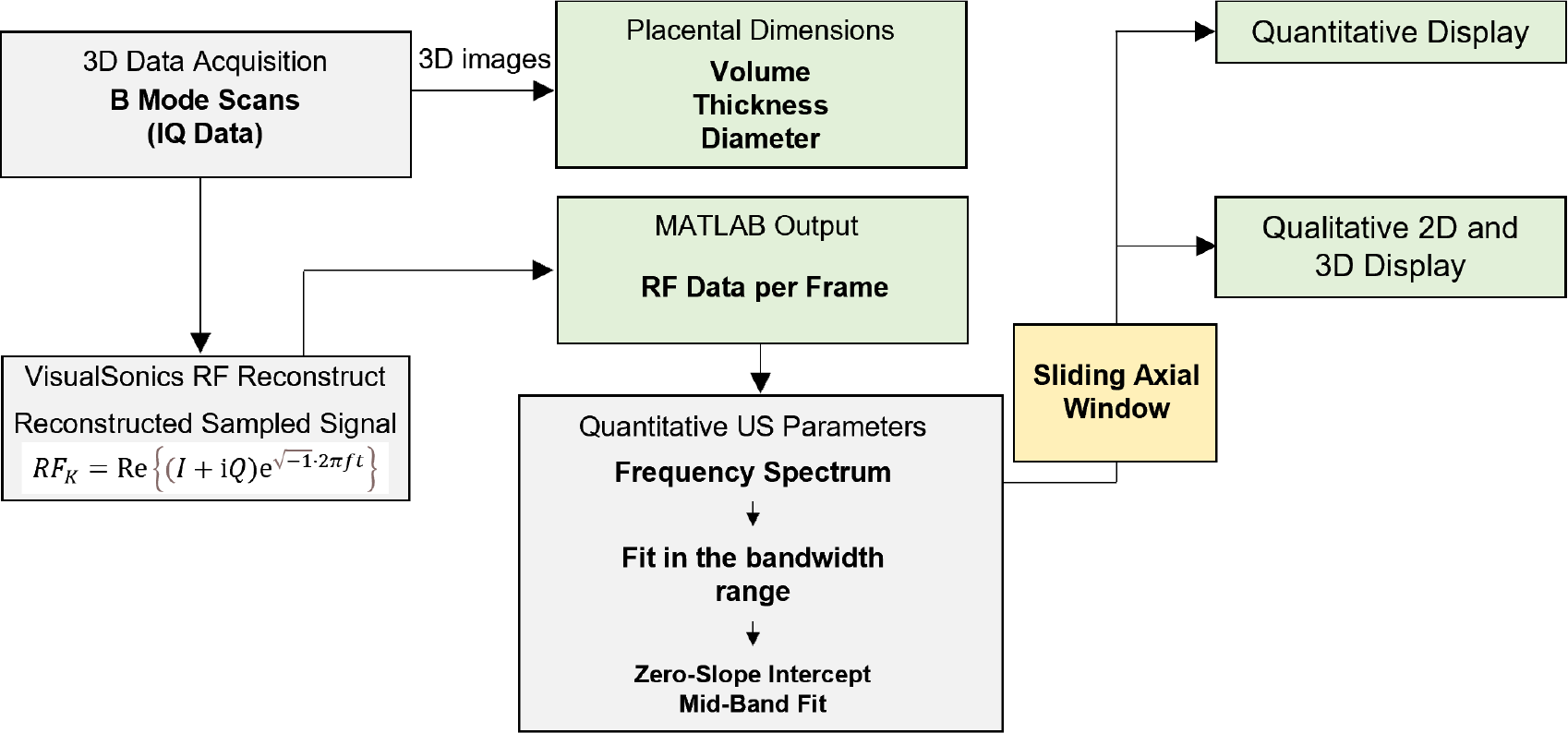
Quantitative ultrasound workflow employed in the study.

### Histology

In this study, cryo-sections of tumor/placenta tissues (thickness of 10 microns) on glass microscope slides, were obtained utilizing a cryotome. Post fixation with acetone and methanol mixture, hematoxylin and eosin (H&E) staining procedures, as previously described by Xavierselvan et al.^42^ Alizarin Red (AR) stain was performed using a method specified as in Speer et al., except for a 1 h 15 min Alizarin Red treatment time.^43^ For immunofluorescence staining, the cryo-sections were initially subjected to fixation by immersion in a pre-cooled mixture of acetone and methanol at a 1:1 ratio for a duration of 10 minutes. Subsequently, the sections were air-dried for 30 minutes and then subjected to three sequential 5-minute washes with PBS. Following this, a blocking solution with a 1x concentration (Thermo Scientific) was applied to the sections for an incubation period of 1 hour at room temperature. Primary antibodies, namely mouse CD31/PECAM-1 affinity purified polyclonal Ab (R&D Systems Inc) and FITC-Mab (Hydroxyprobe Inc), were administered at an approximate concentration of 10 μg/mL. The incubation of these antibodies was carried out overnight at 4 °C. On the subsequent day, a secondary antibody, specifically donkey anti-goat IgG NL493 affinity purified PAb (R&D Systems Inc), was introduced following washing with PBS. This secondary antibody was allowed to incubate at room temperature for 2 hours. The slides were thoroughly rinsed and sealed utilizing a SlowFade Gold antifade reagent with 4’,6-diamidino-2-phenylindole (DAPI) (Invitrogen, Thermo Fisher Scientific) and coverslips.^44^ The stained slides were imaged at 4x magnification with an EVOS fluorescence imaging system.

### Statistical Analysis

GraphPad Prism (La Jolla, CA, USA) was used to calculate statistical analyses. An unpaired parametric one-sided t-test with 95% significance level was used to compare the oxygenation and QUS parameters across the two placental genotypes respectively. A *p*-value < 0.05 is considered statistically significant.

## Results and Discussion

### 1. KO placentae are smaller in size compared to WT placentae

Figure 3 depicts the statistical comparison between the dimensions of the WT and KO placentae. The top row (Figs. 3a-c) demonstrates the methodology used to calculate the volume, thickness, and diameter of the placenta respectively. Specifically, the collection of 3D volumetric scans (X-Y axes plane is the traditional B-scan imaging plane and Z-axis is the elevational direction) of the *ex vivo* placenta enabled the measurement of placental volume (entire 3D volume), thickness (X-Y plane), and diameter (X-Z plane). As some of the placentae are elliptical, to obtain diameter of the placenta, an average of two diameters was taken as shown in Fig. 3c. The bottom row of Fig. 3 displays the statistical differences between the WT and KO placentae. In concurrence with a previous study that measured placenta thickness through histological assessment, we observed the KO placentae have lower diameter and thickness than the WT placenta in our study.^40^ We also observed that there was no significant difference between the volumes of the two types of placenta. This discrepancy may be due to the age range of the mice used for breeding. In Fig. 3d, mothers from all ages were included and we did not want to exclude any data. However as shown in Supplementary Fig. 1a, b, there is a significant difference in placental volume and thickness depending on the age of the pregnant mouse. This age-dependent variation emphasizes the need to not only study models that express ectopic calcification, but also to investigate the impact of maternal age on placental health. The inconsistency in placental size is not unique to our study and suggests that other metrics are needed to determine placental dysfunction. ^45-47^

**Figure 3:**
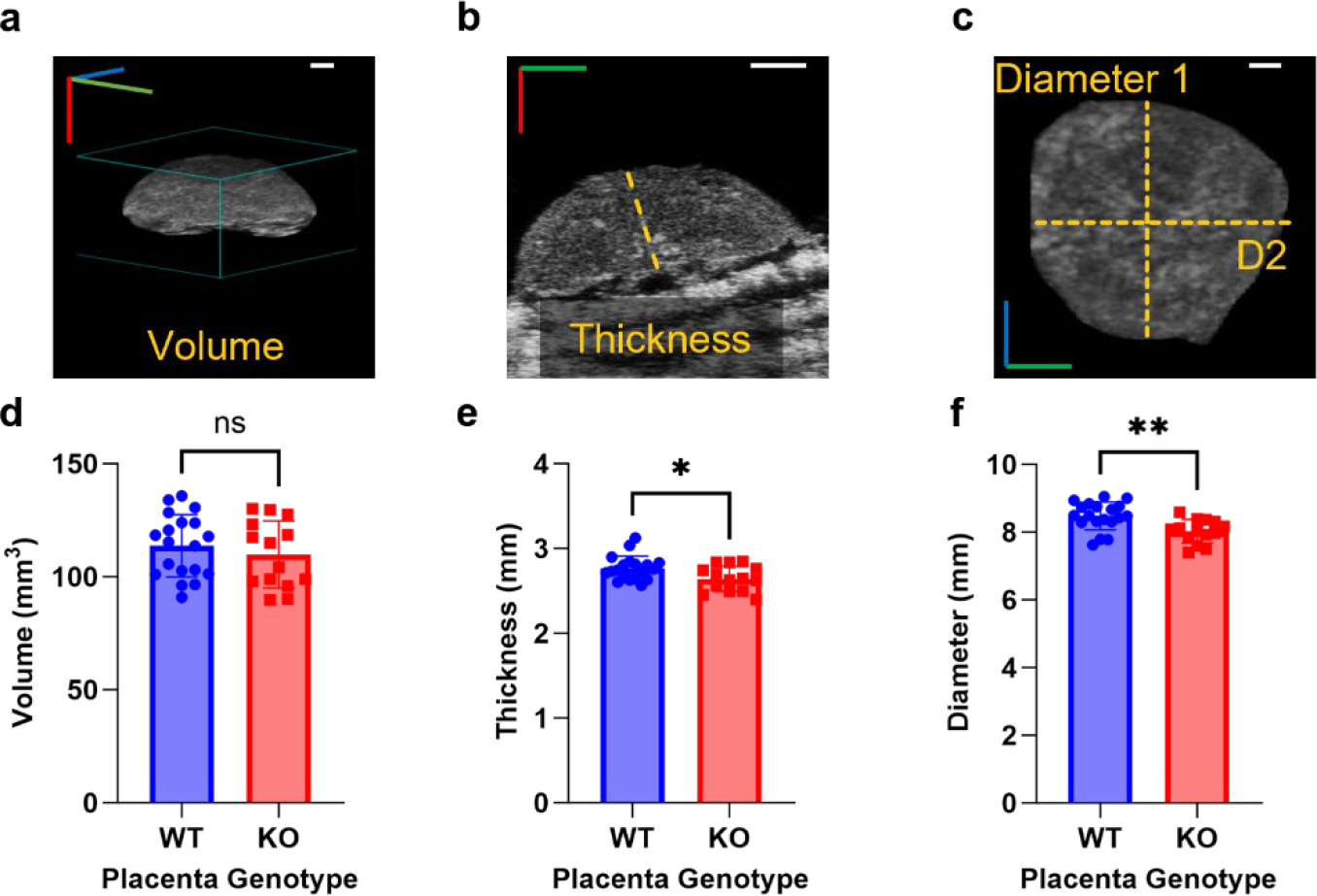
*Ex vivo* US measurements of collected the placentae. **A)** The volume of KO and WT placentae. **B)** The thickness of the middle frame of the placentae (*p*-value <0.05). **C)** The average diameter of the collected placentae (*p*-value <0.05). **D-F)** Examples of the measurements done on a single placenta. Each graph was tested with an unpaired two tailed t-test. The green axis is the x-direction, red is y-direction, and blue is z-direction. Scale bar = 1 mm. N_WT_= 19 placentae; N_KO_= 14 placentae.

### 2. KO placentae have higher calcification and hypoxia compared to WT placentae

Distinct regions of the murine placenta are shown in Fig 4a. The placenta is made up of maternal decidua, the fetal-derived junctional zone, and the meshed maternal-fetal vascular interface (labyrinth).^48^ Figures 5b and 5c depict immunofluorescence staining in which blue is a stain for cell nuclei (DAPI), red indicates the endothelial cell marker CD31, and green is pimonidazole which is used to assess hypoxia. The CD31 signal (Figs. 4b) identified maternal blood vessels in the decidua and revealed the highly vascularized regions of the labyrinth. Both the WT and KO placentae have similar vascular density observed qualitatively through CD31 staining, but the blood vessels in the KO placenta experience more hypoxic conditions represented by increased pimonidazole signal in the KO compared to the WT placenta (shown by the white arrows Figs. 4b insets). Importantly, the hypoxia corresponds with the Alizarin Red staining in similar locations of the labyrinth from a WT and KO placenta seen in Figs. 4c and d. The calcified red nodules depict that the KO placenta has more calcified lesions when compared to the WT placenta, as previously reported.^40^ After validating the Slc20a2 model through histological analysis and identifying increased hypoxia, we developed tools to differentiate between the two types of placenta *in vivo* using quantitative ultrasound and photoacoustic imaging (QUSPAI).

**Figure 4:**
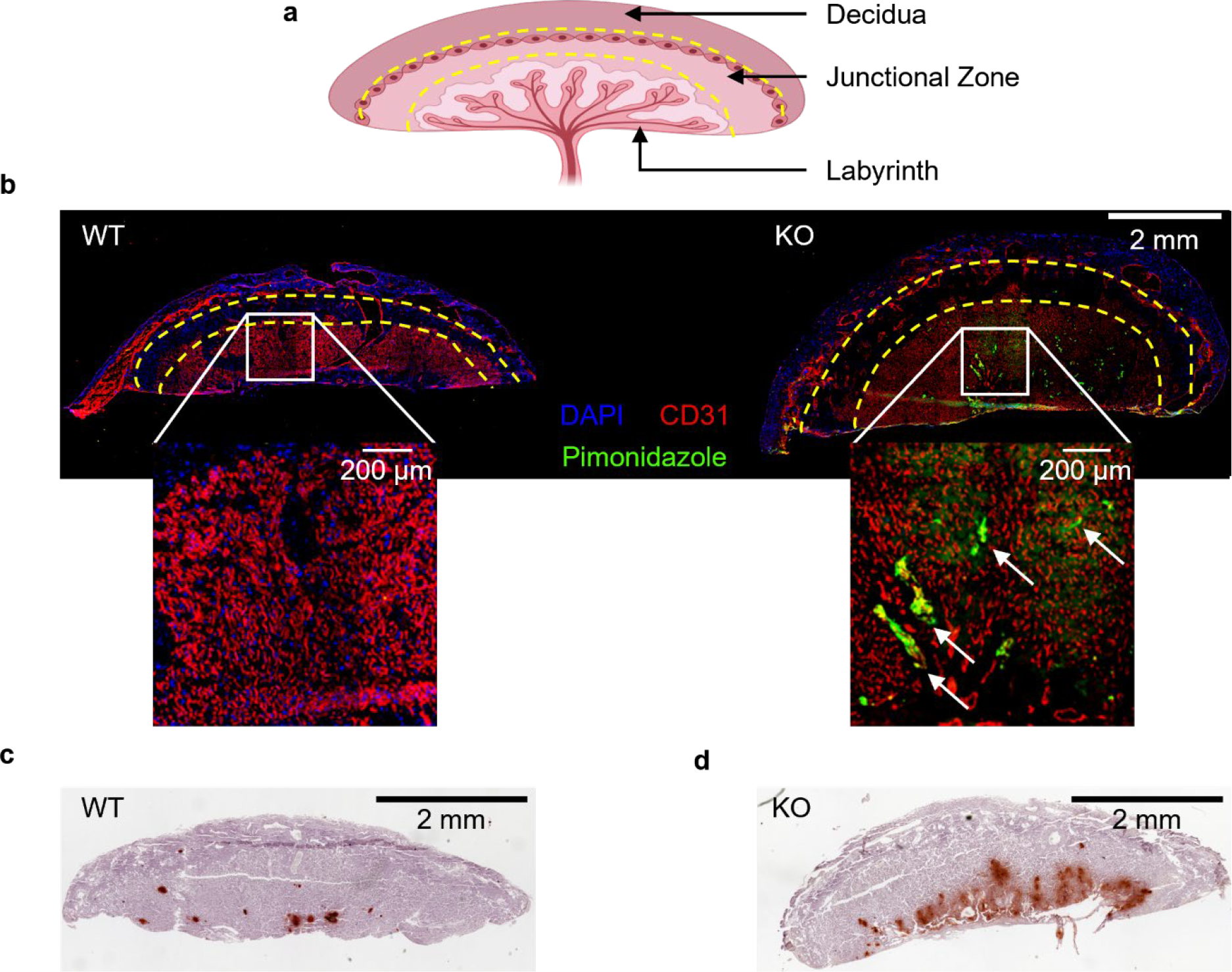
**a)** Schematic of a murine placenta with three distinct regions: decidua, junctional zone, and labyrinth. **b)** Immunofluorescence staining. Blue is DAPI, red is CD31, and green is pimonidazole. All figures are collected *in vivo* on embryonic day E18.5. **c, d)** AR-stained slides of a WT and KO placenta. Areas with stronger brown-red stains in the image depict the calcified regions in the KO placenta.

**Figure 5:**
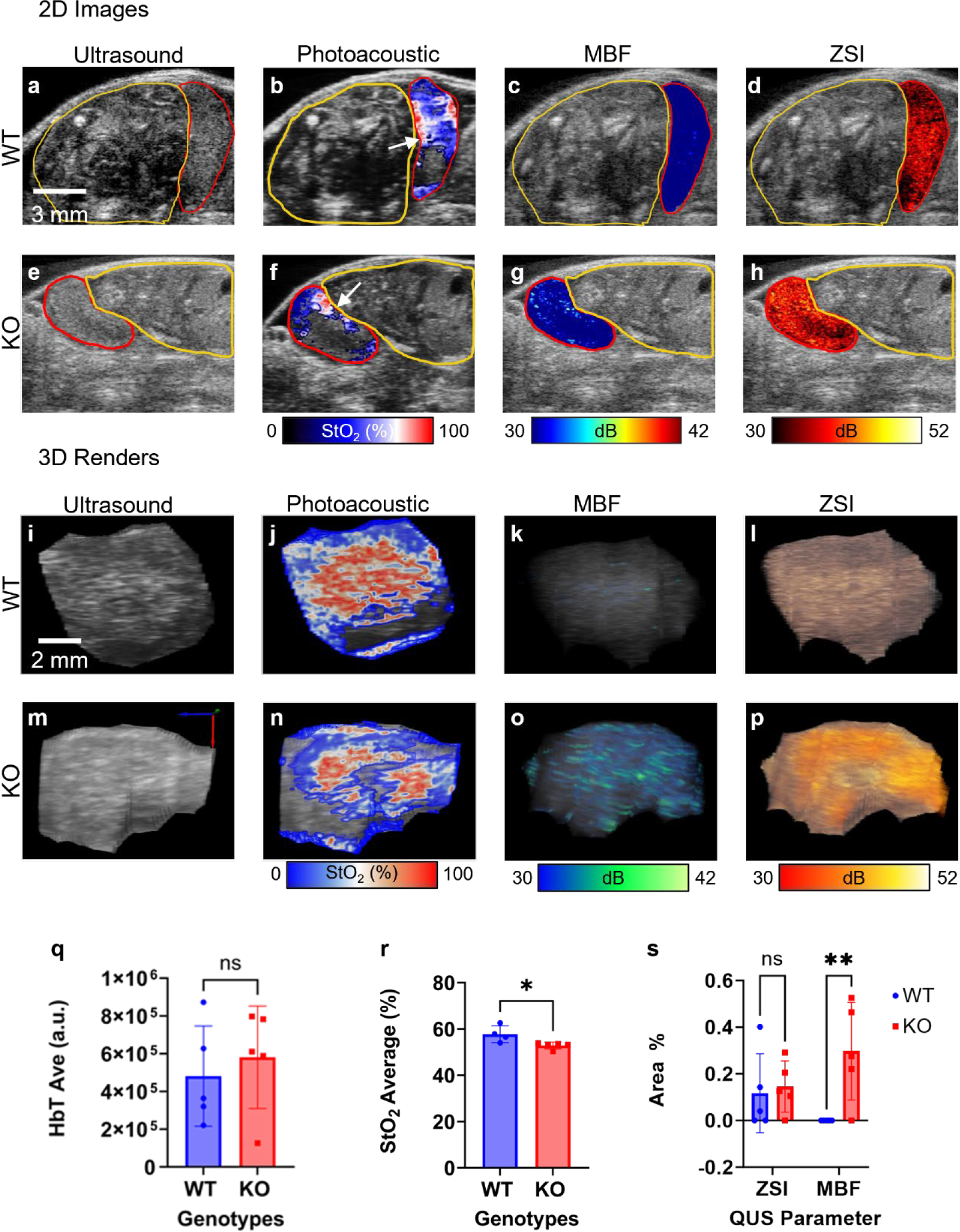
**a-h)** *In vivo* US images, PA StO_2_ map, and QUS maps from two placental scans WT and KO. The first column is grey scale images US, and the second column is PA StO_2_ images. Areas of red indicate oxygenated regions and areas of blue area indicate hypoxic regions. The third and fourth columns are mid-band fit and zero-slope intercept maps. **i-p)** An increased density of the MBF and ZSI peaks can be observed from the KO to the WT placenta, especially along the chorionic plate of the KO. **q)** It was indicated that total hemoglobin content is not significantly different between genotypes (*p*-value <0.05). **r)** We determined that the WT placenta StO_2_ average was statistically significant from the KO placenta (*p*-value <0.05). **s)** Comparing the QUS parameters MBF and ZSI, separately, indicates that ZSI is not a parameter that is as significant in determining *in vivo* identification (*p*-value <0.05). All tests were unpaired parametric one tailed t-test. All figures are collected *in vivo* embryonic day E18.5; mid-band fit color bar range is 30 to 42 dB; and zero-slope intercept color bar axis is 40 to 52 dB. N_WT_= 4 placentae; N_KO_= 5 placentae.

### 3. Multi-parametric imaging reveals strain-specific placental characteristics

QUSPAI allows functional and structural imaging of the placenta *in vivo*. Figure 5 depicts 2D and 3D multi-parametric maps from a representative WT and KO placenta where Figs. 5a-h depict a cross section that is parallel to the sagittal plane of the placenta, while Figs. 5i-p shows a 3D rendering of the placentae cut on the transverse plane. The first column depicts a grayscale US image in 2D (Figs. 5a and 5e) or 3D (Figs. 5i and 5m). US imaging provides anatomical and structural information of the mouse abdomen and is used to determine the region of interest (ROI), the placenta in this case. The embryo and the placenta are outlined by the yellow and red ROIs respectively in the 2D images (Figs. 5a-h). In the 3D images (Figs. 5i-p), we segmented the placenta in the 2D images and only show the 3D reconstruction of the placenta. The orientation of the 3D axes for both the placentae are also shown in Supplementary V1 and V2 where Figs. 5i-l and m-p are shown at the same cut plane.

The second column of Fig. 5 introduces the complementary PA images obtained at the same cross section as the US images. The PA images show the placental StO_2_ values where red pixels indicate highly oxygenated regions, blue pixels indicate hypoxic regions, and black pixels were replaced to be transparent to indicate no signal. A comparison of the WT and KO placentae reveals several key differences. Within the 2D images the labyrinth regions identified by white arrows are densely packed with blood vessels and hence have high StO_2_ values. However, the differences between the genotypes are not as apparent in the 2D StO_2_ images and it is key to observe the 3D USPA imaging images as shown in Figs. 5j and 5n. Specifically, in the 3D images we show underneath the placenta and the total length of the labyrinth, where clearly the KO placenta has lower StO_2_ values overall than WT placenta. This is in agreement with the data seen in Figs. 4b and c, where the KO placenta has similar vessel density observed in the labyrinth area, but higher pimonidazole (hypoxia) stain. Quantification of the PA images, post compensating for fluence differences due to the location of the placentae, yielded statistically significant differences between the WT and KO for only the StO_2_ measurements (Fig. 5r) but not the total HbT values (Fig. 5q) as expected. This indicates that there is a decreased oxygenation in the KO placentae compared to the WT placentae, while the overall blood vessel density remains the same in both types of placentae.

The third column of Fig. 5 represents the MBF parameter, where areas of yellow or red indicate a strong signal of acoustic impedance, increased effective scatterer size, and concentration. The fourth column of Fig. 5 represents the ZSI parameter, where areas of yellow and white indicate increased effective scatterer size and concentration. These parameters are the same in the 3D images, however the color maps are different due to the use of different software for rendering the images. QUS and PAI imaging modalities are taken sequentially, and the mouse’s breathing during live imaging can adjust the position of the embryo and placenta in between scans. As seen in Figs. 5c and d and Figs. 5g and h there is a visual difference between the WT and KO QUS parameters in each placenta. In agreement with histology, Fig. 4d and e, where the AR stain shows dense and punctate calcifications in the labyrinth region, the MBF images show the punctate high MBF signals in the labyrinth region and near the decidua. The ZSI images are less concentrated into one region of the placenta and instead can be seen throughout both WT and KO. Where these two parameters overlap there is an increased acoustic impedance and an increased concentration of acoustic scatterers as anticipated because the calcified regions of the placenta are expected to be denser and stiffer compared to the other regions of the placenta. The MBF and ZSI parameters were compared with area percentage, the ratio of the number of QUS pixels above the threshold, and the total number of US pixels present in the frame between WT and KO placentae using a representative QUS midframe, shown in Fig. 5s. *In vivo*, only the MBF parameter provides a statistical difference between the genotypes. This may be due to a larger increase in acoustic impedance, effective scatterer, and/or concentration of scatters combined, compared to just the effective scatterer, and concentration of scatters combined^17, 18^.

Within the placenta, the decidua contains the centralized areas of maternal spiral arteries, and the junctional zone is a throughput to the labyrinth, which is a mesh of fetal and maternal vascular exchanges. The labyrinth is a critical region of the placenta when deciphering placenta dysfunction. In this particular model of placental dysfunction, Slc20a2^-/-^, there are calcifications that occur at the dense vasculature of the labyrinth as seen in histology Fig. 4d. Additionally, hypoxia was observed extensively around these vascular structures in the labyrinth region of the placenta as seen in Fig. 4b. Identifying these regions non-invasively *in vivo* was possible only through the unique combination of QUSPAI. Though the images do not have microscopic resolution, the meso-scale images were quantified to show that the StO_2_ and MBF metrics are statistically different between the WT and KO placentae, pointing out the salient features of QUSPAI.

### 4. Ex vivo QUS maps reveal higher calcifications in KO placentae compared to WT placentae

*Ex vivo* QUS imaging was performed to validate our *in vivo* results that higher ectopic calcification was present in the KO placenta. We utilized a high frequency 40 MHz transducer that allowed obtaining images at higher resolution and thereby a closer investigation of the calcification striations. In Fig. 6a, a visual representation of US B-scan images (top row), MBF (middle row), and ZSI (bottom row) are shown for the two genotypes investigated. The US B-scan images do not showcase major visual differences between the placentae. However, there is an increased density of the MBF pixels that can be observed from the WT and KO placentae, very similar to *in vivo* images demonstrated in Figs 5c, d, g, and h. Similar differences in the ZSI pixels can also be observed, unlike in the *in vivo* images. Fig. 6b showcases quantified QUS data, where a significant difference between WT and KO in the ZSI and MBF parameters *ex vivo* was observed. Supplementary videos V3 show case B-scans across the length of the placenta where clearly the MBF and ZSI parameters match similar striations to the AR stains in Fig. 4d and e.

**Figure 6:**
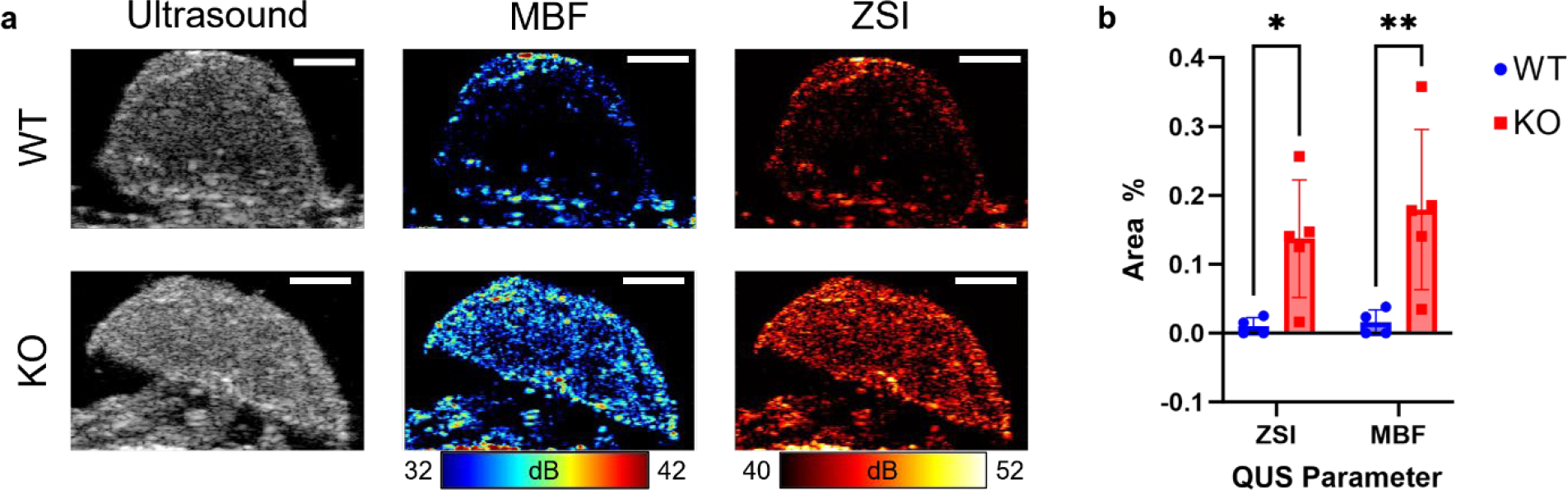
**a)** US images (top), MBF maps (middle), and ZSI maps (bottom) for both genotypes: WT and KO. All figures are collected *ex vivo* embryonic day E18.5; scale bar = 2 mm; mid-band fit color bar range is 32 to 42 dB; and zero-slope intercept color bar axis is 40 to 52 dB. **b)** An unpaired parametric one tailed t-test on both parameters determined that KO placentae have a higher ZSI and MBF ratios compared to the WT placentae (*p*-value <0.1 and *p*-value < 0.05). N_WT_= 4 placentae; N_KO_= 5 placentae.

Hypoxia is a component of early placental development and is essential for a healthy pregnancy, however mid-to-late-stage hypoxia can have harmful effects.^49, 50^ Overall, we found that in the observed late stage E18.5 KO placentae there are sparse oxygenated regions around the labyrinth and increased regions of MBF and ZSI as compared to the WT placentae. The MBF concentrates near the labyrinth indicating an increase in effective scatterer size, acoustic impedance, and concentration of acoustic scatterers. The overlay of QUS parameters and hypoxic regions suggests that abnormal structures in the placenta are related to reduced function of the placenta. These findings suggest that QUSPAI is a valuable tool for identifying and monitoring placental abnormalities in KO mice. The 3D imaging allows for these parametric maps to be gauged at the entire organ level, i.e., parameters can be quantified for the entire placenta, instead of just a representative 2D frame. The 3D imaging of the placenta also enables users to perform quantification of specific regions of interest within the placenta. As previously mentioned, normal B-mode US images had a high variation in between independent observers when grading the calcifications of placentae.^13^ As observed in our study, and echoed in oncology research, QUS has the ability to characterize lesions and reduce the variation in common clinical diagnoses.^26, 51-53^ In essence, 3D QUS imaging can become an indispensable clinical imaging modality to enhance our knowledge of placental development, function and related malignancies.

Translation of the QUSPAI method to the clinic necessitates certain modifications. Firstly, clinical transducers designed for obstetric purposes typically operate in the 4 – 7 MHz range for transabdominal and 5 – 9 MHz range for transvaginal transducers.^54^ While in murine models, we have successfully distinguished MBF and ZSI parameters in healthy and diseased placentae, it’s important to acknowledge that our preclinical approach utilizes high-frequency transducers (different from the clinically available transducers) with limited penetration depth. The clinical transducers can penetrate deeper into the abdomen, albeit at the cost of reduced resolution. Nevertheless, the larger size of placental calcifications in human subjects renders them suitable for imaging and analysis using a similar methodology. Consequently, further exploration of utilizing QUS for evaluation of calcifications in human placenta is warranted. Secondly, pertaining to PA imaging, light delivery to the placenta *in vivo* is a challenge. Murine models have multiple placentae and often these are in superficial locations that enable easy access for USPAI imaging. Conversely in humans, the placental placement is deeper within the abdomen, obscured from light penetration. A potential solution involves employing a customized light delivery system alongside either a transvaginal or transabdominal transducer, akin to methodologies successfully used for imaging ovarian tissues^55, 56 57^. With these developments, QUSPAI can become an invaluable tool in clinics but nevertheless is an extremely valuable tool in studying various mouse models of placental dysfunction.

## Conclusion and Future Work

Understanding *in vivo* ectopic placental calcification is essential for detecting high-risk pregnancies, as it is associated with aging, obesity, and the pregnancy average onset occurring later with the improvement in medicine^58^. Additional preclinical studies are needed to monitor functional and structural parameters over time and at earlier time points. This would help identify when placental insufficiency beings to impact fetal growth. Future work would include compartmental analysis of QUSPAI data, i.e., 3D imaging and demarcating the specific regions such as the labyrinth or the crown of the placenta could also provide insights into the health of the placenta, as the ectopic calcifications tend to develop at the center of placenta compared to lateral regions. The combination of quantitative ultrasound and PA imaging is promising for the diagnosis and monitoring of placental dysfunction. This study demonstrated the feasibility of using QUSPAI to differentiate between healthy and diseased placentae in a preclinical model. While the study has its strengths, future studies are needed to identify the most informative regions of the placenta to image, investigate the impact of maternal age on placental health, and validate QUSPAI in a clinical setting to assess its ability to predict placental insufficiency and adverse pregnancy outcomes. QUSPAI has the potential to revolutionize the management of high-risk pregnancies by providing real-time information about placental health.

## Acknowledgements

The authors would like to acknowledge funds from NIH S10 0D026844 (Mallidi), Tufts University School of Engineering (Mallidi), Tufts University Data Intensive Studies Center (Mallidi), and the GEM Fellowship Program (Mallidi), NIH K99/R00HD090198 (Wallingford), AHA 19CDA34660038 (Wallingford), the Forbes Family (Wallingford), the Susan Saltonstall Foundation (Wallingford), the Herbert J. Levine Foundation (Wallingford) and the Tufts Medical Center Mother Infant Research Institute (Wallingford). The authors would also like to thank Tomoko Kaneko-Tarui at the Tufts Medical Center and Maureen Kehler at the Tufts University Comparative Medicine Services.

## Supplementary Information

**Supplementary Figure 1:**
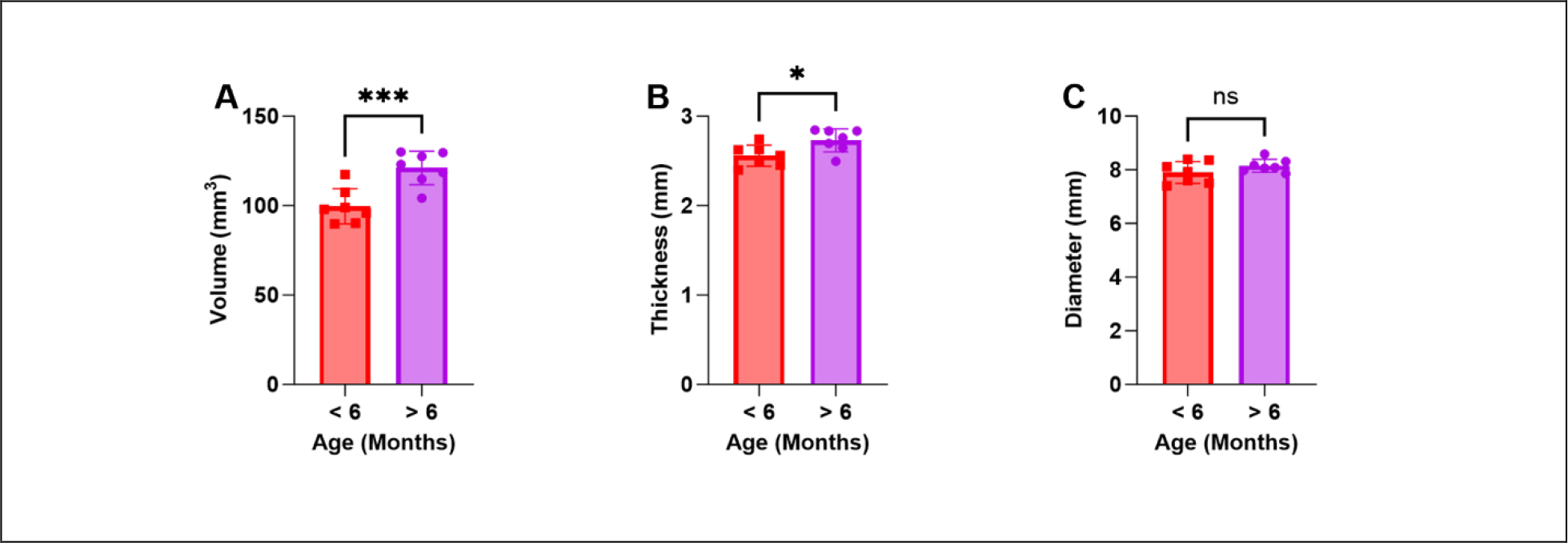
Red are KO placentae collected from mice younger than six months. Purple are KO placentae collected from mice older than six months. An unpaired parametric one tailed t-test indicated the KO placentae that were collected from mice older than six months had larger diameter **(A)**and greater thickness **(B)**. There was no significant difference in the average diameter **(C)** of the placentae from mothers of different age group (unpaired parametric one tailed t-test). N_KO_= 7 placentae for both time frames. * (*p*-value <0.05) *** (*p*-value <0.001)

### Supplementary Videos 1, 2, and 3

Video S1: Video indicating the orientation of the 3D axes for the WT placenta and the imaging plane shown in Figure 5i-l.

Video S2: Video indicating the orientation of the 3D axes for the KO placenta and the imaging plane shown in Figure 5m-p.

Video S3: Movie of the B-scans across the length of the two placental types. The MBF and ZSI parameters are both shown here where the MBF is displayed with a jet colormap (range: 32 to 42 dB) and the ZSI is displayed with a hot colormap (range: 40 to 52 dB).

